# Protein Arginine Methylation of the Translation Initiation Factor eIF1A Increases Usage of a Near-cognate Start Codon

**DOI:** 10.1101/2024.08.16.608280

**Authors:** Rebecca Wegman, Michael Langberg, Richoo B. Davis, Xiaozhuo Liu, Minkui Luo, Michael C. Yu, Sarah E. Walker

**Author notes:** Address correspondence to: M.L, M.C.Y., and S.E.W., Minkui Luo, Chemical Biology Program, Memorial Sloan Kettering Cancer Center, 1275 York Avenue, New York, NY. 10065, Sarah E. Walker, Department of Biological Sciences, State University of New York at Buffalo, 109 Cooke Hall, Buffalo, NY. 14260, Michael C. Yu, Department of Biological Sciences, State University of New York at Buffalo 109 Cooke Hall, Buffalo, NY. 14260. These authors contributed equally to this work (Rebecca Wegman, Michael Langberg, Minkui Luo).

## Abstract

Protein arginine methylation has emerged as a key post-translational modification responsible for many facets of eukaryotic gene expression. To better understand the extent of this modification in cellular pathways, we carried out bioorthogonal methylation profiling in *Saccharomyces cerevisiae* to comprehensively identify the *in vivo* substrates of the major yeast protein arginine methyltransferase Hmt1. Gene ontology analysis of candidate substrates revealed an enrichment of proteins involved in the process of translation. We verified one such factor, eIF1A, by *in vitro* methylation. Three sites on eIF1A were found to be responsible for its methylation: R13, R14, and R62, with varied capacity by which each site contributed to the overall methylation capacity *in vitro*. To determine the role of methylation in eIF1A function, we used a battery of arginine-to-alanine substitution mutants to evaluate translation fidelity in these mutants. Our data show that substitution mutants at R13 and R14 in the N-terminal tail improved the fidelity of start codon recognition in an initiation fidelity assay. Overall, our data suggest that Hmt1-mediated methylation of eIF1A fine-tunes the fidelity of start codon recognition for proper translation initiation.

## Introduction

Protein arginine methylation has emerged as a key regulator in protein function and is catalyzed by a family of evolutionarily conserved enzymes called protein arginine methyltransferases (PRMTs) (1–3). PRMTs utilize S-adenosyl-*L*-methionine (SAM) as the methyl donor in methyltransferase reaction, and their SAM-binding pocket is formed by motifs I, post-I, and the THW loop. These enzymes can be classified based on the forms of methylarginine generated: type I PRMTs catalyze the formation of either monomethylarginine (MMA) or asymmetric dimethylarginine (aDMA), whereas type II PRMTs catalyze MMA or symmetric dimethylarginines (sDMA). Hmt1, the budding yeast homologue of mammalian PRMT1, is the major type I PRMT whose activity is responsible for the formation of 89% of aDMA and 66% of MMA *in vivo* (4, 5). Hmt1 preferentially methylates arginine residues in the context of RXR/RG/RGG motifs (6), and previous large-scale and targeted studies have revealed only ∼40 Hmt1 *in vivo* substrates, of which RNA-binding proteins represent the largest class of proteins identified (7–9). The number of *in vivo* Hmt1 substrates found contrasts largely with a recent finding in mammalian PRMT1, where hundreds of substrates have been identified via proteomic profiling (10). Such disparity hinted at the likelihood where many more Hmt1 *in vivo* substrates have yet to be identified.

Previous attempts at identifying the full scope of Hmt1 substrates utilized either proteomic shotgun mass spectrometry or affinity purification of Hmt1-associated proteins (7–9). These attempts were insufficient at identifying large numbers of novel substrates because they were either 1) trying to identify protein-specific methylation marks in the context of a total proteome, a difficult task due to sensitivity level, or 2) capturing proteins that were associated with Hmt1 but not necessarily being modified by the enzyme. More importantly, these approaches lack the resolving power to effectively identify weak or transiently associated substrates of Hmt1. To address these caveats, bioorthogonal profiling technology has emerged as a robust method for identifying *in vivo* substrates of specific protein methyltransferases (PMTs) (11). In this technology, the desired PMT is altered to accommodate SAM analogues (e.g. 4-azido-but-2-enyl-SAM or Ab-SAM) that are bulkier than native SAM. Since these analogues are expected to be more specific for tailor-engineered PMT variants rather than the native enzymes (12), implementation of this approach will allow for the transfer of a bioorthogonal azido moiety to substrates specific to the engineered PMT (12). Such tailor-engineered PMT is subjected to the cellular proteome where the transfer from SAM analogues to substrates can be carried out, thereby allowing substrate proteins to be labeled in the process. The resulting labeled proteome is then subjected to biotinylation with a “Click-iT” biotin-DIBO alkyne, followed by streptavidin pulldown and identification using mass spectrometry (12). Indeed, this approach has been successfully implemented to profile substrates of several classes of PMTs (13).

Post-translational modifications (PTMs) such as arginine methylation have previously been identified in proteins involved in eukaryotic translation but only one example of methylation of the core translation initiation machinery has been characterized to date (14, 15). During eukaryotic translation initiation, a start codon is recognized through a scanning mechanism by a preinitiation complex (PIC). This PIC is formed when a ternary complex (TC), including the initiator tRNA (Met-tRNA_i_^Met^) and eIF2-GTP, binds to the 40S ribosomal subunit with other essential initiation factors. This complex binds near the m^7^G-capped 5’-end of the mRNA and moves along the untranslated region using the anticodon of the initiator tRNA to identify a start codon, typically the first AUG in a good sequence context. The conformation of the 40S and the initiator tRNA are controlled by several eukaryotic initiation factors (eIFs) including eIF1A, which modulates these PIC conformations. The binding of eIF1A to the 40S subunit shoulder is achieved by the OB fold of this factor, while the conformation of the PIC is modulated through interactions of its C- and N-terminal tails, to stabilize TC in either the P_out_ or P_in_ state, as well as a 40S open or closed conformation. While the tRNA and mRNA are loaded and the PIC is scanning an mRNA, the 40S occupies an open conformation with the TC in a P_out_ state that allows for movement along, and examination of the mRNA to recognize a start codon (16). Once a start codon basepairs with tRNA, the tRNA is accommodated in a P_in_ state, which clashes with eIF1 leading to reorganization of the network of interactions to stabilize a closed conformation of the PIC, arrest scanning, and allow 60S subunit joining and subsequent elongation (16, 17). Hence, regulating this balance between the open/P_out_ and closed/P_in_ conformations is critical for accurate start codon recognition and the overall translation process. One key mechanism by which this balance could be regulated is via PTM of eIFs. Indeed, a role for phosphorylation of eIFs in regulation of early translation initiation events has previously been described (18, 19). However, much less is known about how arginine methylation contributes to the function of eIFs in the process of translation.

In the work presented here, we utilized the bioorthogonal profiling approach to comprehensively identify *in vivo* substrates of Hmt1-mediated methylation. Gene ontology analysis of the identified Hmt1 *in vivo* substrates reveals the highest enrichment in gene expression, RNA metabolic process, and RNA binding categories. Furthermore, we have identified eIF1A as a candidate *in vivo* substrate from our profiling data and verified its capacity to act as a substrate for Hmt1 *in vitro*. We generated substitution mutants at residues and identified R13, R14, and R62 as methylarginines in eIF1A. The capacity by which each of the methylated arginine residues contribute to the overall methylation of eIF1A *in vitro* was determined. The functional role for these methylarginines in translation initiation was tested via a translation start codon fidelity assay and we found substitution mutants of R13 and R14 greatly improved fidelity and inhibit usage of a near-cognate start codon. These results suggesting that methylation of R13 and R14 promote the open/P_out_ conformation of the PIC to allow for optimal start codon initiation.

## Experimental Procedures

### Yeast Strains and Growth Conditions

All yeast strains used are listed in Table S1. Cells were grown at 30°C on YPD medium (1% yeast extract, 2% Bacto peptone, 2% w/v D-glucose) unless otherwise stated. Gene deletions and integration of epitope tag was performed as previously described (20, 21). All plasmids used are listed in Table S2 and all primers used are listed in Table S3. To generate the yeast strain (MYY 2894) that overexpresses modified Hmt1 (Hmt1-M36G), site-directed mutagenesis was carried out on a copy of *HMT1* gene cloned into the NotI site on the plasmid pSP400 where its expression is under the control of entire *ADH1* promoter and terminator. This resulted in a plasmid that contains the M36G substitution, which is then used to generate the Hmt1-M36G mutant yeast strain by inserting *HMT1-M36G* sequence flanked by a *URA3* sequence into the endogenous *HMT1* locus in a *hmt1Δ* strain (MYY 432). To generate the yeast strain (MYY 1298) that expresses a version of Hmt1 that is catalytically inactive (Hmt1-G68R), we cloned the gene that expresses this version of Hmt1 (from pPS1760) into the NotI site on the pSP400 plasmid. Both the Hmt1-M36G-containing or Hmt1-G68R-containing plasmid were linearized with StuI prior to use in yeast transformation into MYY 432, resulting in either MYY 2894 (for Hmt1-M36G) or MYY 1298 (for Hmt1-G68R). Yeast strains harboring eIF1A point mutations were selected for on synthetic complete medium (SC) lacking leucine (Sunrise Science, cat#1304-030 and cat#1650-250). Resident eIF1A p3392 was replaced through plasmid shuffling by plating on media containing 5-fluoroorotic acid (GoldBio, cat# F230-25), as described previously (16, 22). Derivatives containing *sui5^-^* (p4281, eIF5^G31R^) were selected on SC lacking leucine and tryptophan (Sunrise Science, cat#1317-030). Medium used for start codon fidelity assays were generated with synthetic dextrose minimal medium (Sunrise Science, cat#1739-500 and cat#1650-250) with respective amino acids added depending on desired medium.

### Scintillation Assay for Recombinant Hmt1 Activity

The *in vitro* methylation reaction testing Hmt1 and Hmt1-M36G was carried out as previously described (23) with the following differences: for Hmt1 and Hmt1-M36G, 2.5µg of the enzyme was incubated with 4µg of recombinantly purified Npl3 and 0.75µM of tritiated *S*-adenosyl-L-methionine ([methyl-^3^H]-SAM) at room temperature overnight in a 20µl total reaction volume. The reaction was quenched by blotting reaction mixtures onto squares of P81 cation-exchange filter paper (GE Healthcare) in three 6µl aliquots. Filter paper was neutralized by washing in bicarbonate and excess SAM was removed before cutting the filter paper into individual squares, mixed with Ultima Gold scintillation solution (Perkin-Elmer) and analyzed with a Perkin Elmer Tri-Carb 2910 TR Liquid Scintillation Analyzer. Data were analyzed in GraphPad prism and presented as mean ± standard error of the mean. Each data point represents a technical triplicate of three reads from a single methylation reaction.

### Synthesis of SAM Analogs

Hey-SAM and Pob-SAM were synthesized as previously reported using reagents purchased from Sigma-Aldrich and used without further purification (12). Hey-SAM and Pob-SAM were purified by HPLC (Waters 600 Controller HPLC with an XBridge™ 103 Prep C18 5μm OBD™ 19×150mm column), lyophilized, and stored in 0.01% trifluoroacetic acid before use.

### Labeling of Yeast Cell Lysates with Synthetic SAM Analogs

Single colonies of either *HMT1-M36G* or *hmtlΔ* were grown in YPD media (10 ml total culture volume for in-gel fluorescence and mass shift assays. For mass spectrometry profiling experiments, colonies were grown overnight then diluted into 500 ml YPD and grown either overnight (for stationary phase experiments) or until reaching an OD_600_ of 0.8-1.0 (for log phase experiments). Cells were then pelleted by centrifugation and resuspended in ten times cell pellet volume of methyltransferase reaction buffer (50 mM HEPES pH 8, 100 mM KCl, 15 mM MgCl_2_, 10% glycerol) supplemented with 1mM PMSF, 5 mM TCEP, and protease inhibitor cocktail (Roche). Cells were lysed by addition of 1.5g acid-washed sterile glass beads (Sigma) per gram pelleted cells and 5 rounds of vortexing (1 min each), with 1 min incubation on ice between each round of vortexing. Lysates were separated from glass beads by piercing the bottom of containers and drained by low-speed centrifugation. Lysates were then cleared by centrifugation at 16.1k RCF for 20 mins and its protein concentration quantified. Equal total masses of protein were used for each condition of interest (typically 50-100 μg total protein for in-gel fluorescence experiments and 10 mg total protein for proteomic experiments) and incubated with 100 μM cofactor (Hey-SAM or Pob-SAM) and 100 nM recombinant MTAN protein (to prevent product inhibition of methyltransferase activity) overnight. Following incubation, the reaction was quenched by protein precipitation either through addition of 3:2:1 methanol:water:chloroform followed by centrifugation for 10 mins at 16.1k RCF or addition of 25x reaction volume of ice cold methanol and overnight precipitation at −80°C. Regardless of precipitation method, the precipitated proteins were washed two times with cold methanol followed by centrifugation to isolate a pellet of SAM analog-labeled proteins, which is allowed to air dry prior to copper (I) catalyzed alkyne-azide cycloaddition (CuAAC) reaction.

### Copper (I) catalyzed alkyne-azide cycloaddition (CuAAC) reaction

To prepare for the click cocktail, 1 mM CuSO_4_ and 2 mM BTTP ligand (Albert Einstein College of Medicine Chemical Synthesis Core Facility) was premixed followed by addition of 2.5 mM sodium ascorbate (Sigma-Aldrich) to reduce the Cu^2+^ to its active catalyst Cu^+^ state. 250 μM azide reagent is then mixed with the active catalyst to yield our 4-component click cocktail. For in-gel fluorescence experiments, TAMRA-azide (Life Technologies) was used as the azide reagent and for biotin-pulldown experiments, cleavable Diazo Biotin-Azide (Click Chemistry Tool) was used. Dried SAM analog labeled protein pellets were completely resuspended in click buffer (50 mM TEA, 150 mM NaCl, 2% SDS), followed by the addition of the click cocktail before the reaction incubated under constant mixing in dark for 90 minutes. Reaction was quenched by protein precipitation as described above and protein pellets were washed twice with cold methanol and air dried.

### In-gel Fluorescence of SAM Analog-Labeled Proteins

Dried protein pellets were resuspended in loading buffer (40 mM Tris pH 6.8, 70 mM SDS, 10 mM EDTA, 10% glycerol, 10% β-ME) without dye to avoid interference with fluorescence signal and separated by SDS-PAGE. SDS-PAGE gels were fixed in 40% methanol, 10% acetic acid overnight to wash out free TAMRA dye and reduce background. After fixing overnight the gel was rinsed in double distilled water to rehydrate and scanned for fluorescence signal using the TAMRA channel on a Typhoon TRIO variable mode imager (Amersham Bioscience). After fluorescence scanning the gel was stained with coomassie blue to confirm equal protein loading. All steps prior to scanning the gel on the Typhoon imager were performed with samples covered to prevent loss of fluorescence signal through photobleaching

### In vitro methylation assays with autoradiographic readouts

The *in vitro* methylation assay was carried out as previously described (23). For Hmt1 and Hmt1-M36G, 2.5µg of the enzyme was incubated with 4µg of recombinant Npl3 and 0.75µM of tritiated S-adenosyl-L-methionine ([methyl-^3^H]-SAM) at room temperature overnight. The methylation reactions were resolved on an SDS-PAGE gel, followed by transfer to a PVDF membrane. The membrane was stained with ponceau S followed by either fluorography or autoradiography.

### Purification of intein-tagged eIF1A

Protein purification was completed as described previously (24), except that 300 mL of LB supplemented with carbenicillin (100 mg/ml) and chloramphenicol (34 mg/ml) was used, and protein was concentrated and stored following elution from the chitin column.

### Translation start-codon fidelity assay

The histidine growth assay for translation start-codon fidelity was carried out as described previously (22). Yeast strains harboring indicated *TIFΔ* (eIF1A) alleles (25) were grown in liquid medium overnight at 30°C. The cultures were diluted to an OD_600_ of 0.5 prior to use. Ten-fold dilutions were plated on selection plates in the presence (0.3mM His, +His) or absence (0.003mM His, -His) of histidine. Plates were incubated at 30°C for 8 days. The same procedure was carried out to compare *HMT1* with *hmt1Δ* mutant strains.

### Experimental Design and Statistical Rationale

For Hmt1-M36G TMT mass spectrometry analysis, triplicate biological samples were carried out. For *in vitro* methylation, at least three technical replicates were performed, including the stated controls. For the translation fidelity assays, at least three biological replicates with at least two technical replicates were carried out in the experiment, including the stated controls.

## Results

### Engineered Hmt1-M36G alkylates native Hmt1 substrate Npl3

To comprehensively profile Hmt1 *in vivo* substrates using the bioorthogonal profiling technique, we first engineered Hmt1 to perform an alkylation reaction of native substrates in the presence of a synthetic cofactor. This approach was previously carried out for PRMT1 (26) and and PRMT3 (13), two human homologues of Hmt1. In the two cases, a conserved methionine (M48 of PRMT1 and M233 of PRMT3) was mutated to glycine and the resulting engineered PRMTs were successfully used in bioorthogonal profiling of human cell lines (26). Upon comparing the amino acid sequences and structural alignments of PRMT1, PRMT3 and Hmt1, it was found that the conserved methionine aligns with the methionine at position 36 of Hmt1 (Fig. 1A). Using the same rationale, we substituted the methionine at position 36 of Hmt1 with glycine (Hmt1-M36G) and utilized a scintillation assay to test Hmt1-M36G’s capacity in methylating Npl3, a robust native substrate of Hmt1 (Fig. 1B) (27, 28). Our data showed that Hmt1-M36G displayed significantly decreased activity toward radiolabeled SAM when compared to Hmt1 (Fig. 1B, comparing Hmt1+Npl3 bar to Hmt1-M36G + Npl3 bar). This decrease was anticipated and desired for efficient SAM analog labeling by Hmt1-M36G in cell lysates, during which the synthetic cofactors must compete with endogenous SAM. The observed difference provides an opportunity to use synthetic SAM analogs to achieve higher Hmt1-mediated labeling of cellular proteins, which can then be tracked in subsequent proteomic work-flow.

**Figure 1.**
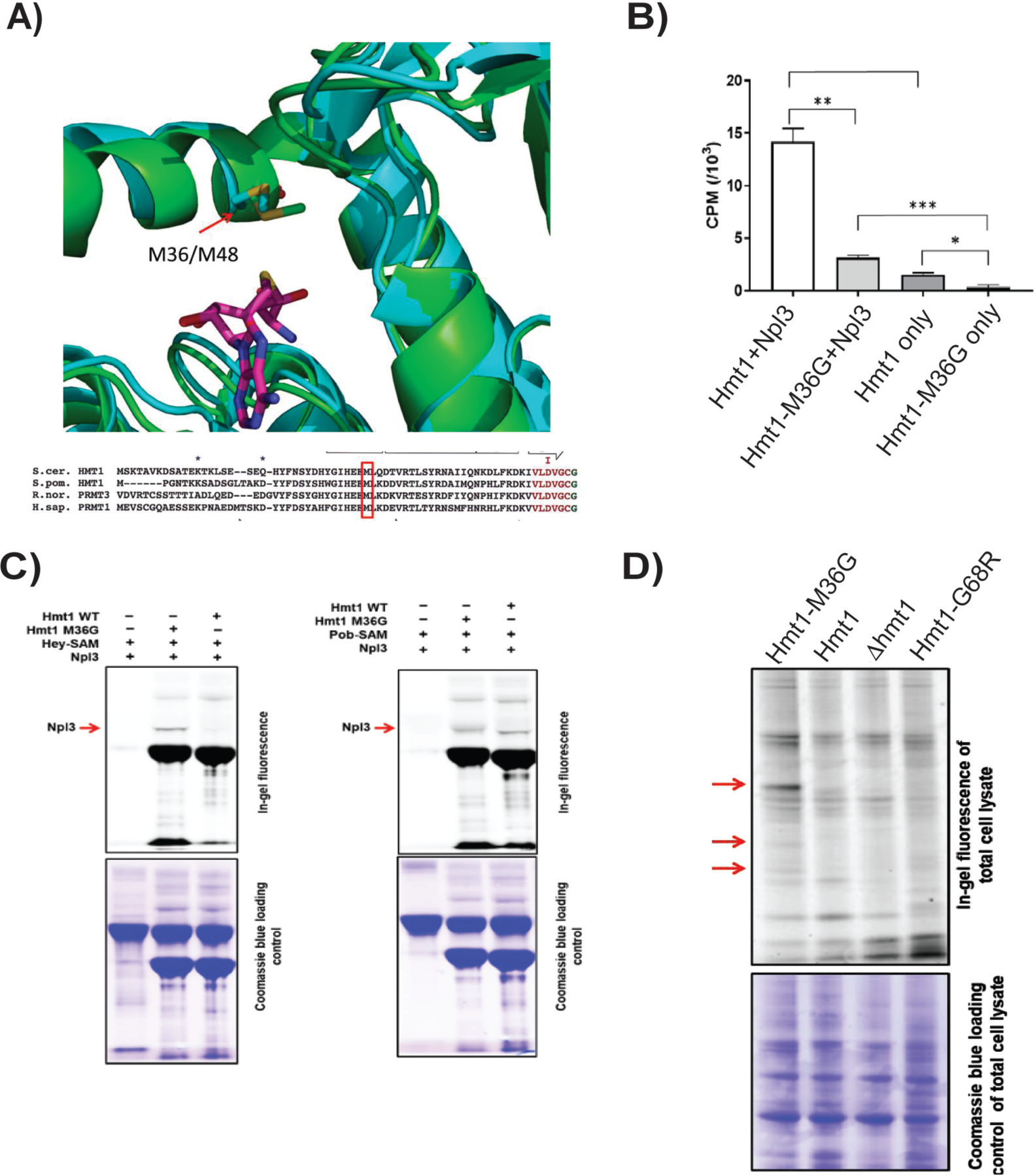
Tailored engineered yeast major protein arginine methyltransferase Hmt1-M36G preferentially utilizes *S-*adenosyl-L-methionine analogues. **a)** Sequence and structural alignment of *S. cerevisae* Hmt1 with *S. pombe, R. norvegicus,* and *H. sapian* PRMT1. The boxed amino acid at position 48 represents a highly conserved methionine residue in PRMT1, which corresponds to residue at position 36 in Hmt1 in both the three-dimensional structure of the two proteins (ribbon structure on the top panel) and the primary amino acid sequence (sequence alignment on the lower panel). **b)** Labeling of Npl3 by Hmt1 or Hmt1-M36G. Recombinantly purified Hmt1 or Hmt1-M36G was incubated with recombinantly purified Npl3 overnight in the presence of tritiated SAM followed by activity measurement by scintillation assay. The result plotted represents technical triplicates of three reads from a single methylation reaction. *P-*values were calculated using a two-tailed t-test, with *<=0.05, **<=0.001, and ***<=0.001. **c)** Specific *in vitro* labeling of Npl3 by Hmt1-M36G and Hey-SAM. The ability of Npl3 being labeled by Hmt1 or Hmt1-M36G using either Hey-SAM or Pob-SAM as a cofactor was measured by an in-gel fluorescence assay. The arrow denotes the band corresponding to labeled Npl3 by Hmt1 or Hmt1-M36G. **d)** In-gel fluorescence labeling assay was carried out using the following yeast lysates harvested in log phase: Hmt1-M36G, wild-type (parental), *hmtlΔ*, and Hmt1-G68R. The lysates were subjected to CuAAC in the presence of Hey-SAM and TAMRA-azide, followed by SDS-PAGE. The gels were subjected to analysis for fluorescence labeling (top panel) and coomassie staining (bottom) for loading control. Red arrows indicate bands that appear only in the lysates from Hmt1-M36G.

We next determined which of the following synthetic cofactors, Hey-SAM or Pob-SAM, would be effectively utilized by Hmt1-M36G to modify Npl3. Using an in-gel fluorescence labeling assay (Fig. 1C), we demonstrated that Hmt1-M36G is able to utilize either Hey-SAM or Pob-SAM to modify Npl3, with Pob-SAM being more effective for a higher level of alkylation based on the observed mass shift. In contrast, Hmt1 was unable to modify Npl3 at all using Hey-SAM (Fig. 1C). Similar to Hmt1-M36G, Hmt1 appeared to be capable of modifying Npl3 using Pob-SAM to certain degree (Fig. 1C). This observation led us to conclude that using Hey-SAM in our bioorthogonal profiling assay would provide a lower background and therefore more specific labeling of cell lysates when compared to using Pob-SAM (Fig. 1C).

We then compared the ability of Hmt1-M36G versus that of Hmt1 in broad labeling of proteins in yeast cell lysates in the presence of Hey-SAM. To this end, we prepared yeast cell lysates from Hmt1, Hmt1-M36G, Hmt1-G68R (a previously published catalytically inactive mutant of Hmt1 (29)), or *hmtlΔ* cells grown to log phase and subjected each lysate to CuAAC in the presence of Hey-SAM and TAMRA-azide, followed by analysis for fluorescence labeling (Fig. 1D). Our data revealed several reproducible bands appearing only in lysates from Hmt1-M36G (Fig. 1D, red arrows). In parallel, we carried out the same experiment using lysates harvested from the stationary phase and observed the same reproducible bands in the Hmt1-M36G lane only, although a lower background was observed with the log-phase when compared to that of the stationary phase (data not shown). Hence, our data demonstrate the feasibility of selective labeling by Hmt1-M36G in the context of cell lysates isolated from log-phase grown cells.

### Large-scale Substrate Profiling of Hmt1-M36G Using Tandem Mass Tag

We employed a quantitative mass spectrometry technique incorporating both negative and positive control groups, enabling the identification of non-specific factors associated and enrichment of true targets. This technique, termed tandem mass tagging (TMT), makes use of easily identifiable “barcodes” appended to peptides at the end of the processing for mass spectrometry analysis (Fig. 2A). To carry out this proteomic workflow, triplicate samples of matched Hmt1-M36G and *hmtlΔ* lysates were prepared from log-phase cells, and the resulting lysates were treated with Hey-SAM. Subsequently, TMT mass spectrometry analysis was performed on our triplicate Hmt1-M36G and *hmtlΔ* sample pairs. We identified 872 proteins present in all three of our biological samples, with 540 showing enrichment in our Hmt1-M36G data set (Fig. 2B). A curated list was generated comprising all proteins with a Hmt1-M36G/*hmtlΔ* ratio greater than 1 and a *p-*value ≥ 0.05, including proteins with known annotated biological functions and excluding those with unknown roles (Table S4). Among the top 50 hits, 21 of the 41 previously validated Hmt1 substrates were enriched in the Hmt1-M36G samples, while no previously validated substrates were enriched in our *hmtlΔ* control samples (Table S4). To gain insight into the pathways and biological functions in which Hmt1 may be involved, pathway analysis was conducted on our set of 540 enriched proteins, along with an enrichment analysis for gene ontology (GO) terms using yeast-ConsensusPathDB (Fig. 2C). When we calculated the enrichment score and ranked the top ten categories, it was revealed that Hmt1 targets the constellation of pathways involved in translation, such as ribosome, eukaryotic translation initiation, and ribosomal scanning and start codon recognition, as well as RNA metabolism pathways (Fig. 2D). We also compared our study with 88 candidate methylated proteins that were identified by Low et al. using proteomic shotgun approach and the number of overlapped candidates between both studies were only 5 (Fig. 2E).

**Figure 2.**
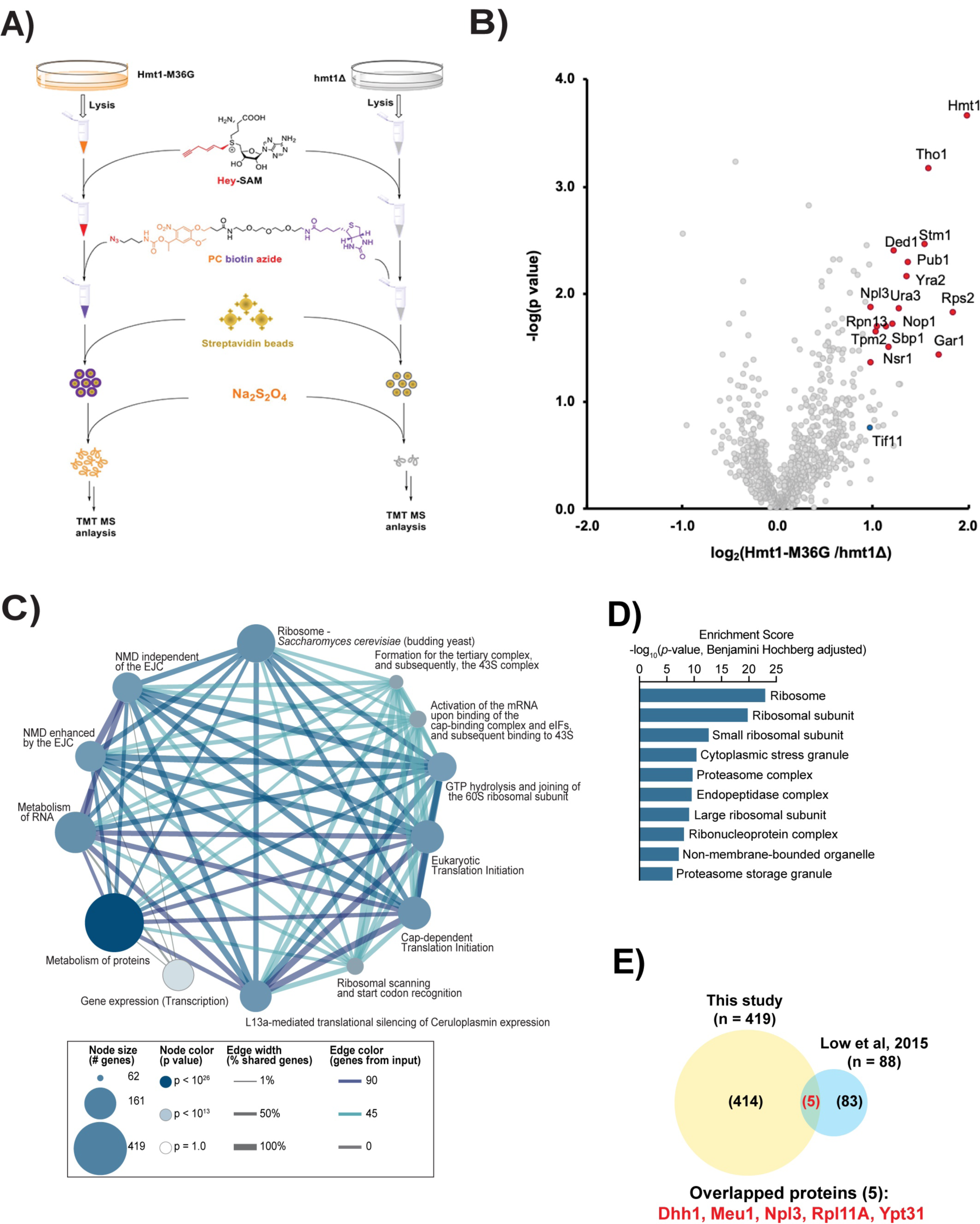
Bioorthogonal profiling of Hmt1-M36G substrates. **a)** Workflow schematic for bioorthogonal profiling of Hmt1 substrates. A yeast strain expressing Hmt1-M36G (MYY 2894) or lacking Hmt1 (MYY 432) was lysed and incubated with Hey-SAM. Following incubation, click chemistry was performed with cleavable biotin azide, and pulled-down peptides were labeled with tandem mass tag (TMT) technology to allow for quantitative mass spectrometry analysis. **b)** Volcano plot of proteins identified via Hmt1-M36G TMT mass spectrometry analysis. Any previously identified Hmt1 substrates with a *p*-value ≤ 0.05 and surpass the median enrichment threshold of 0.115 are labeled with red circles (e.g. Rps2). **c)** Connectivity map of major biological pathways enriched in the TMT mass spectrometry data from Hmt1-M36G substrates. **d)** GO term enrichment analysis of the Hmt1-M36G substrates with calculated score using Benjamini-Hochberg method. Shown are the top ten categories ranked by their scores. **e)** Comparison of Hmt1 candidate substrates identified from this study to those identified by Low et al., 2015. The Venn diagram indicates the number of candidate substrates identified from this study or Low et al., 2015. Of the total number of candidates identified in this study, only those with a mean ratio of 1.1 or above was used to compare with the data from Low et al., 2015. The number of substrates identified from both studies is indicated at the intersection of both circles and listed.

### Translation initiation factor eIF1A is methylated by Hmt1 in vitro

Our profiling data revealed several candidate translation initiation factors modified by Hmt1 *in vivo* (Table S4). One such factor is eIF1A (encoded by the gene *TIF11*), which plays a critical role in ensuring the fidelity of start codon recognition (30). To confirm the methylation of yeast eIF1A by Hmt1, we carried out an *in vitro* methylation assay using recombinantly purified eIF1A (31) (Fig 3A). The fluorograph of the *in vitro* methylation assay exhibited a strong tritium signal at approximately 17 kDa, corresponding to the expected molecular weight of yeast eIF1A (Fig. 3A, left panel, lane eIF1A + Hmt1). The ponceau S stained PVDF membrane also revealed a single band corresponding to the same 17 kDa band observed on the fluorograph (Fig 3A; right panel, lane eIF1A + Hmt1). The appearance of the 17 kDa band in the fluorograph was attributed to Hmt1 activity, as it was absent in the reaction that lacks Hmt1 (Fig 3A; left panel, lane eIF1A – Hmt1). Moreover, Hmt1 activity in the absence of eIF1A did not produce the 17 kDa band (Fig 3A; left panel, enzyme only). Thus, our findings corroborated the results obtained from Hmt1 bioorthogonal profiling, confirming eIF1A as a substrate of Hmt1.

**Figure 3.**
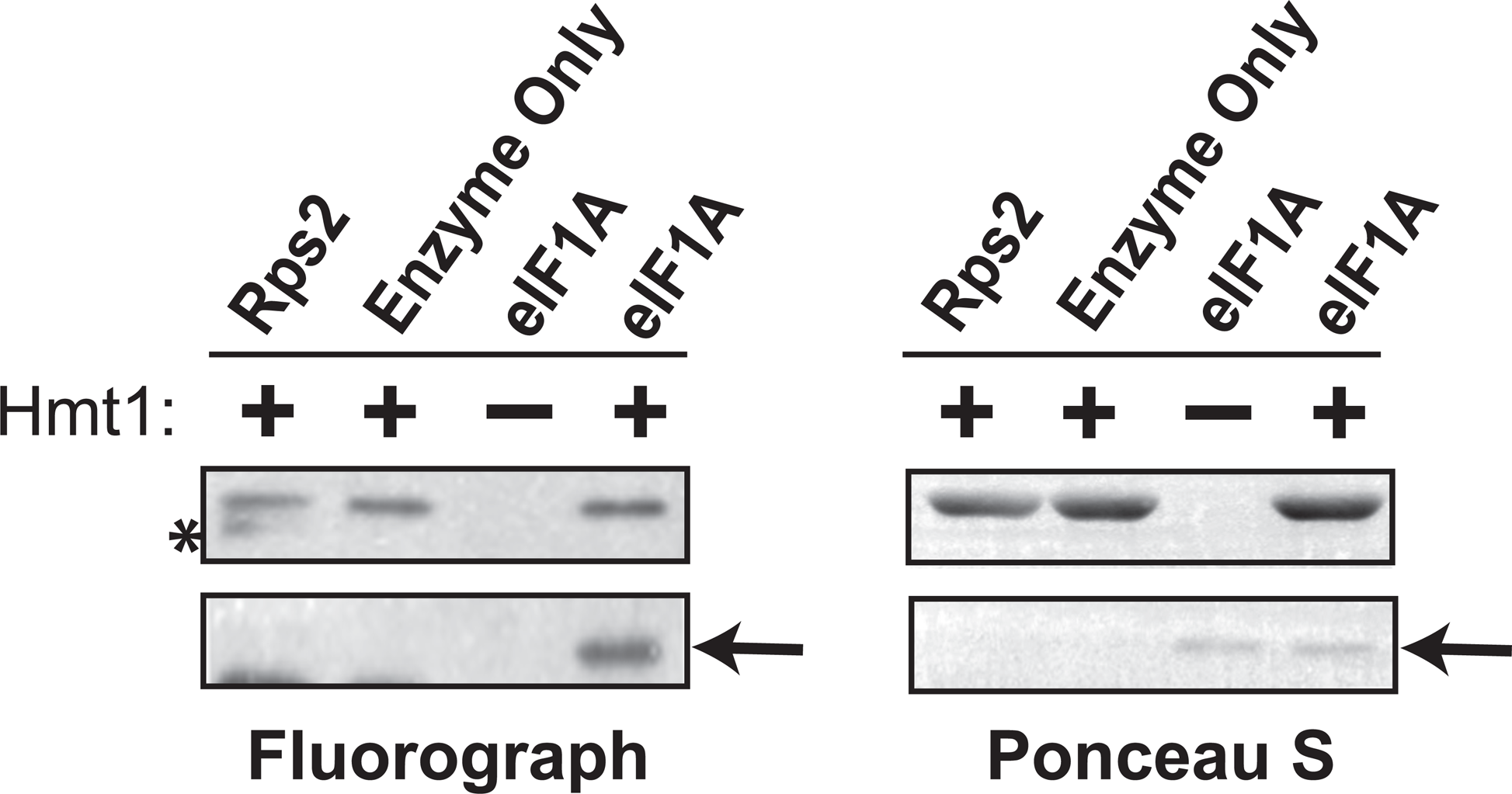
Yeast eIF1A/Tif11 is able to act as an *in vitro* substrate of Hmt1. *In vitro* methylation of intein-tagged wild-type eIF1A/Tif11, following purification from *E. coli*, was carried out using recombinant Hmt1 and [methyl-^3^H]-SAM. The full protein complement in each reaction was resolved on a 4-12% SDS-PAGE and transferred to a PVDF membrane. Methylated eIF1A/Tif11 was visualized by fluorography (arrow), after Ponceau S staining of the membrane to demonstrate protein loading levels (arrow). Recombinant GST-tagged Rps2 served as a positive control (highlighted by asterisk).

### Methylation of eIF1A occurs on arginine residues 13,14 and 62

Amino acid sequence analysis of eIF1A identified three potential sites for Hmt1-mediated arginine methylation: R13, R14, and R62 (Fig. 4A). These sites reside within, or adjacent to, PRMT1/Hmt1-preferred methylation motifs RG/RGG/RXR (32, 33). To assess the methylation potential of each residue by Hmt1, we generated a series of arginine-to-alanine substitution mutants at these positions and subjected them to an *in vitro* methylation assay (Fig. 4A). Among the single substitution mutants, R13A and R14A displayed a noticeable decrease in methylation signal compared to wild-type eIF1A, with R14A showing the most significant impact on methylation signal (Fig. 4B; compare eIF1A^WT^ to eIF1A^R13A^ or eIF1A^R14A^). In the R13,14A double substitution mutants, a greater reduction in overall methylation signal relative to the wild-type was observed compared to the difference between wild-type and the single R14A mutant (Fig. 4B; compare eIF1A^WT^ compared to eIF1A^R14A^ and eIF1A^R13,14A^). These findings suggest that R13 and R14 contribute differentially to eIF1A methylation capacity, with R14 being a critical residue for methylation of eIF1A. Notably, methylation signal was completely abolished in a triple mutant where all three arginine residues were substituted, compared to the R13,14A double mutant (Fig. 4B; eIF1A^R13.14A^ compared to eIF1A^R13,14,62A^), indicating that R62 also contributes to the overall eIF1A methylation signal, albeit to a lesser extent than R13 and R14 individually. Overall, our data demonstrate that residues R13, R14, and R62 are the only arginine residues within eIF1A methylated by Hmt1.

**Figure 4.**
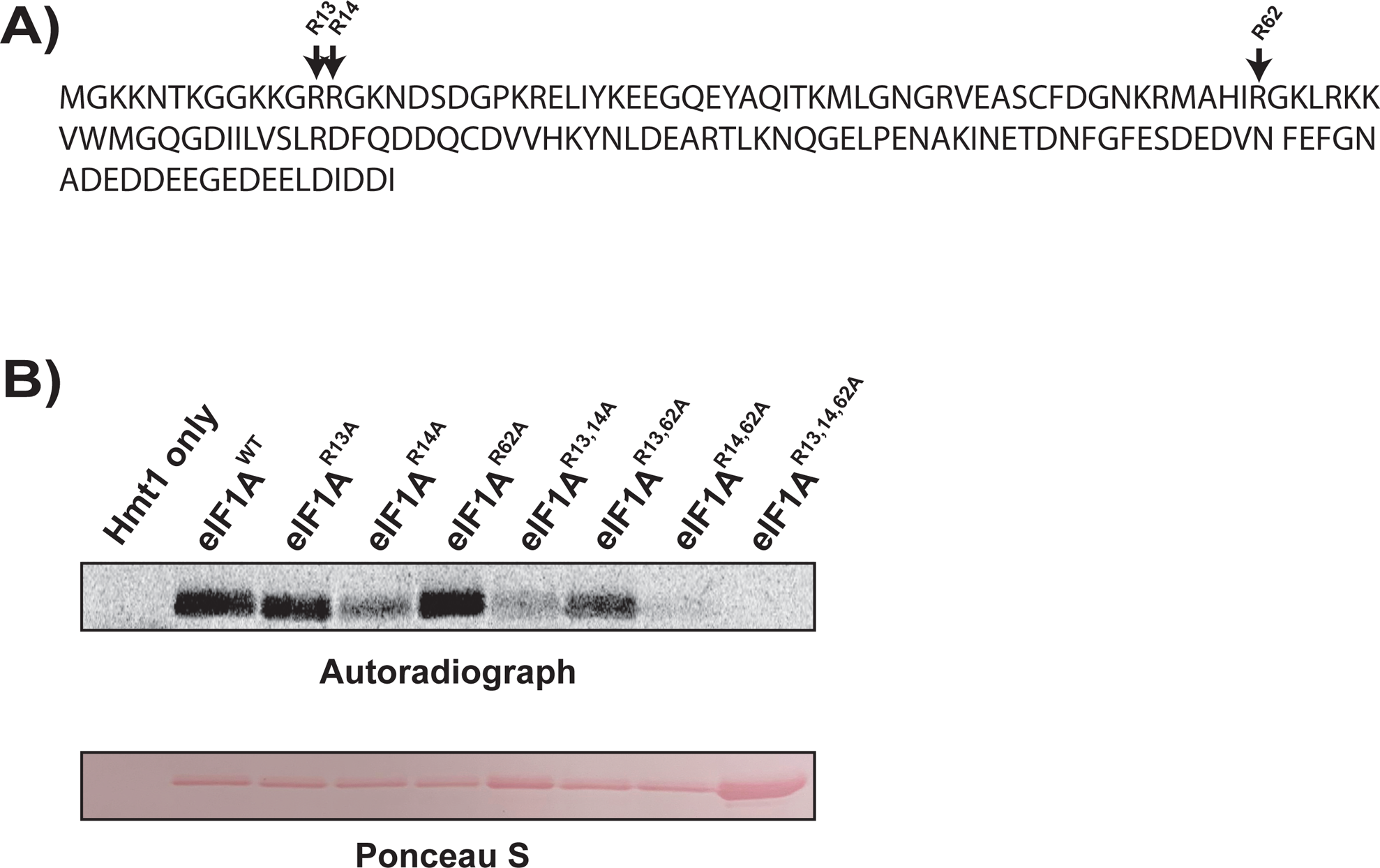
R13, R14, and R62 differentially contribute to the overall methylation capacity of eIF1A/Tif11. **a)** The amino acid sequence of yeast eIF1A/Tif11, with potential methylated arginine sites at positions 13, 14, and 62 denoted by arrow. **b)** Intein-tagged wild-type eIF1A/Tif11 or eIF1A/Tif11 harboring R to A substitution(s) at amino acid positions 13, 14, and 62 were purified from *E. coli* and subjected to an *in vitro* methylation assay using recombinant Hmt1 and [methyl-^3^H] SAM as described in Figure 3. Methylated eIF1A/Tif11 was visualized by autoradiography (top panel), following Ponceau S staining of the membrane to demonstrate protein loading levels (bottom panel).

### Fidelity of start codon recognition is increased in hmt1 loss-of-function mutants in vivo

To investigate how methylation impacts the function of eIF1A, we first generated both *HMT1* and *hmt1Δ* mutants in the *his4-301* yeast strain and used a genetic assay for translation initiation fidelity to determine how Hmt1 affects translation start codon recognition. The *his4-301* yeast strain cannot survive in the absence of added histidine, as the cognate AUG start codon of the *HIS4* gene has been replaced. However, expression of the *HIS4* gene and growth on media lacking sufficient histidine (-His) can be restored if the strain harbors a suppressor of initiation codon mutation (sui^-^) that allows recognition of a near-cognate UUG start site. The sui^-^ class of mutations confers decreased translation initiation fidelity, allowing recognition of a near-cognate start codon and histidine production by promoting the closed/P_in_ state of the PIC (34–38). In contrast, suppressor of sui- (ssu-) mutations have also been isolated that are capable of suppressing the HIS+ phenotypes conferred by sui- mutations. These ssu^-^ mutations enhance fidelity and counteract the growth conferred by sui^-^ mutants on histidine-lacking media by decreasing occupancy of the closed/P_in_ conformation of the PIC and preventing translation of the *HIS4* gene. Numerous sui- and ssu- mutations of eIF1A have been characterized, primarily residing in the C- and N-terminal tails of the protein, respectively (16, 39–42). As Hmt1 methylates an arginine residue in the scanning inhibitory N-terminal tail of eIF1A (Fig. 4), known to influence initiation fidelity (16), we hypothesized that deletion of *HMT1* would confer an ssu^-^ phenotype.

To first determine whether *hmt1Δ* mutants exhibited a sui^-^ phenotype, we compared the growth of *his4-301* strains harboring an empty vector (ev) in either *HMT1* or *hmt1Δ* genetic backgrounds on media with or without histidine. Using a spot assay, we observed that *hmt1Δ* mutants did not confer a sui^-^ phenotype as they were unable to promote growth on histidine-lacking medium (Fig. 5; *hmt1Δ*/ev rows). We next tested if this *hmt1Δ* mutant could suppress a well-characterized sui^-^ mutant, thereby conferring an ssu^-^ phenotype. To carry out this analysis, we compared the growth of *his4-301* strains expressing a *sui5^-^* (eIF5^G31R^) plasmid (43) in either *HMT1* or *hmt1Δ* genetic background on medium lacking histidine. In agreement with previous studies, the *sui5-* mutation conferred a slow growth phenotype in the presence of histidine. However, this phenotype was not further affected by *HMT1*. The *sui5^-^* plasmid facilitated growth of the *his4-301* strain on media lacking histidine compared to the empty vector (Fig. 5; compare *HMT1*/*sui5^-^* to *HMT1*/ev), confirming that the eIF5^G31R^ mutation produced the expected HIS+ sui^-^ phenotype. However, the growth of *hmt1Δ*/*sui5^-^* cells was diminished in the absence of histidine (Fig. 5; compare *HMT1*/*sui5^-^* to *hmt1Δ*/*sui5^-^*). Since the deletion of *HMT1* had no effect on growth in the presence of histidine, these data collectively indicate that *hmt1Δ* mutants conferred an ssu^-^ phenotype, suggesting that the presence of Hmt1 normally facilitates recognition of near-cognate start codons.

**Figure 5.**
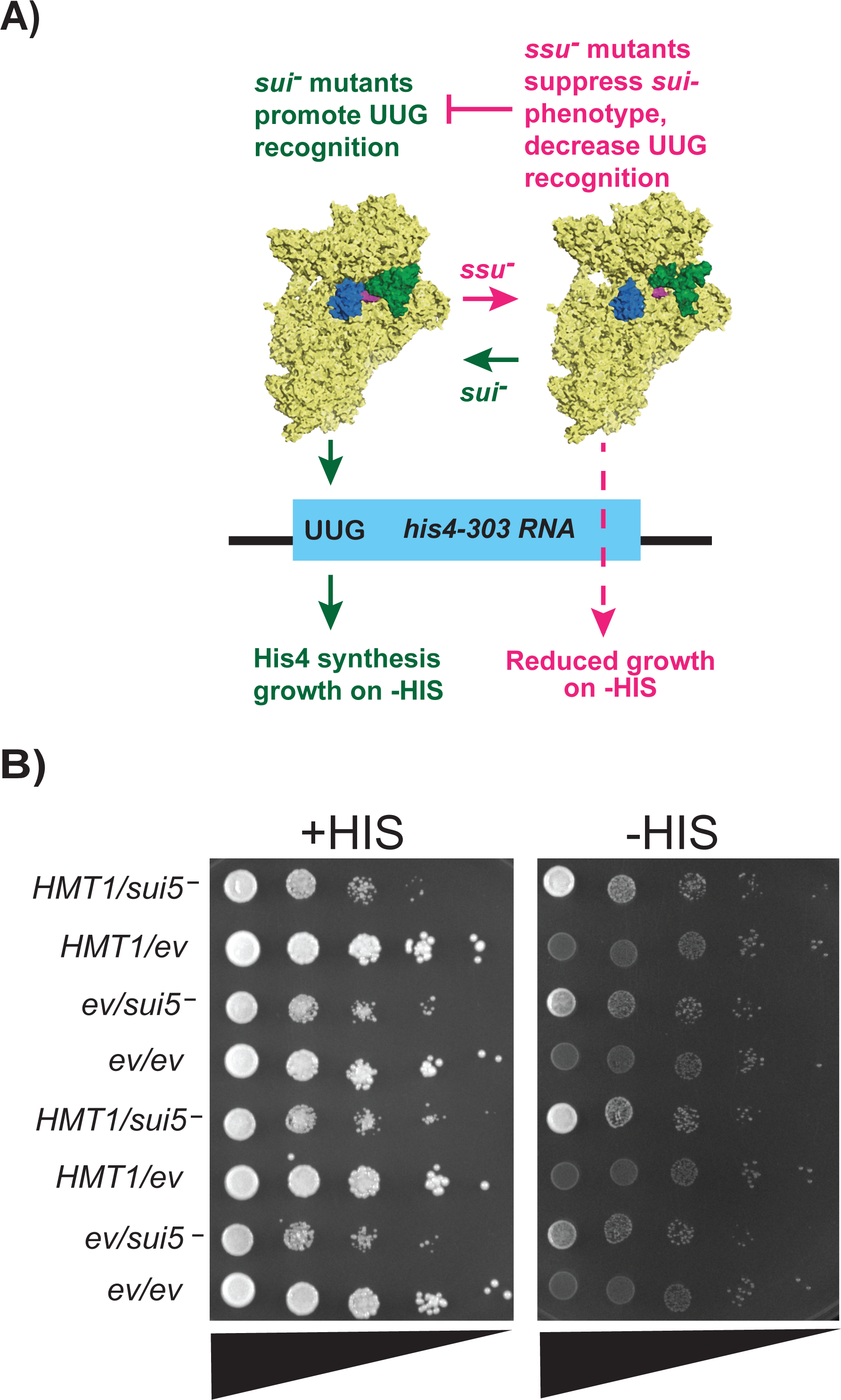
Yeast strain lacking Hmt1 displays partial increase in the usage of proper start codon. a) Overview of the *his4-303* reporter assay used to test for *sui^-^* and *ssu^-^* phenotypes. The *sui^-^* mutations promote a P_in_/closed state of the preinitiation complex, decreasing translation fidelity, and allowing translation of *his4-303* using a near-cognate UUG start codon. This allows histidine biosynthesis and growth on media lacking sufficient concentrations of histidine (-HIS). In contrast, *ssu^-^* substitutions oppose the effects of *sui^-^*mutations to promote the open/P_out_ conformation of the PIC, which prevents translation of the *his4-303* mRNA and prevents growth on -HIS media. Structures of the 43S preinitiation complex were generated in pymol from structures 3J80 and 3J81 from the PDB (38). b) Translation initiation fidelity was assessed in wild-type (*HMT1*) or *hmtlΔ* yeast strains expressing a plasmid harboring indicated mutations in eIF5^G31R^ (MYY 3290 and MYY 3255; indicated by *sui5^-^*) or an empty vector (MYY 3292 and MYY 3257; indicated by ev). A spot assay with ten-fold serial dilution (triangles) was performed using SD medium containing either 0.3 mM histidine (indicated as +HIS) or 0.003 mM histidine (indicated as -HIS). Two biological replicates are shown from the same plate.

### Substitutions in the N-terminal tail of yeast eIF1A increased start codon fidelity in vivo

To investigate whether methylation of arginines at positions 13, 14, or 62 of eIF1A is responsible for the effects of Hmt1 on translation initiation fidelity, we generated a battery of single and combination substitution mutants in which these residues were changed to alanine. We then used the same translation fidelity assay to assess how these mutants affect start codon usage relative to wild-type eIF1A (*TIF11* gene). Our results revealed that none of the single or combination mutants expressing the empty vector (ev) conferred a sui^-^ phenotype (Fig. 6; parts A-C, compare all strains with “ev” in the -HIS panels to *TIF11/sui5-*). Upon examining their abilities to suppress the growth of the sui^-^ mutation *sui5^-^* (eIF5^G31R^), only the single R62A mutant harboring the *sui5^-^* (eIF5^G31R^) plasmid phenocopied wild-type eIF1A expression from the same plasmid (Fig. 5A; compare *TIF11*/*sui5^-^*to *tif11^R62A^*/*sui5^-^*). This observation indicated that the R62A mutant did not confer a ssu^-^ phenotype, suggesting that methylation of that residue does not contribute to translation initiation fidelity. In contrast, either the R13A or R14A single mutant was sufficient to abolish the growth of the sui^-^ mutant on histidine-lacking media, indicating that substitutions of either or both arginine residues in the N-terminal tail, which would prevent their methylation, were sufficient to confer a ssu- phenotype (Fig. 6A; compare *TIF11*/*sui5^-^* to *tif11^R13A^*/*sui5^-^* or *tif11^R14A^*/*sui5^-^*in the -HIS panel). In the double or triple R13, R14, or R62 substitution mutants, incorporation of either the R13A or R14A mutation alone was sufficient to confer ssu- phenotype (Fig. 6B-C; compare *tif11*/*sui5^-^* to mutants in the -HIS panel). When combined with the spot assay data from the *hmt1Δ* mutants (Fig. 5), these findings from single or combination substitution mutants suggest that methylation at the 13^th^ and 14^th^ residues of eIF1A promotes near-cognate start codon usage.

**Figure 6.**
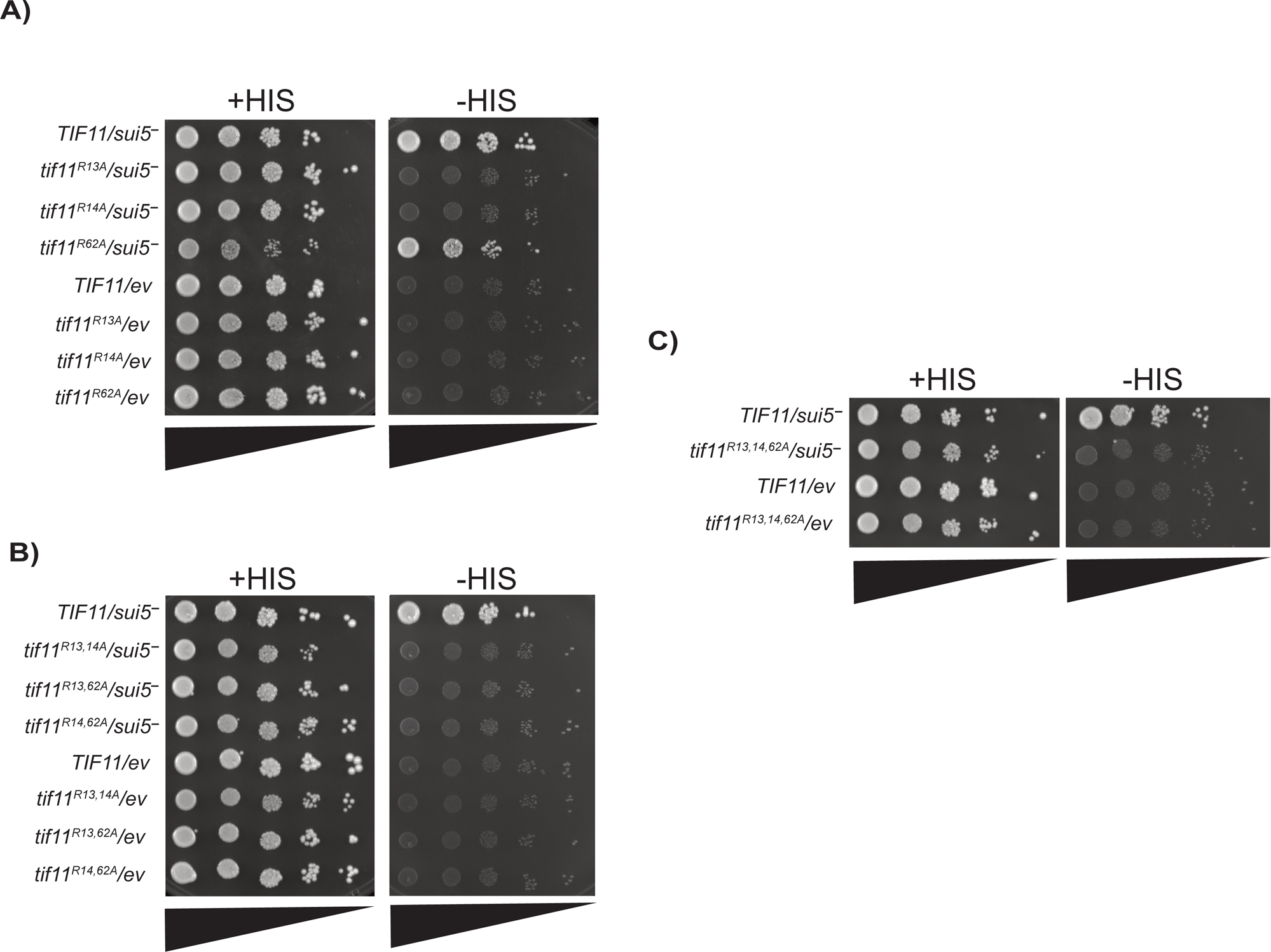
Substitution mutations of eIF1A/Tif11’s R13 and R14, but not R62, increase proper start codon usage when compared to the wild-type. Translation initiation fidelity was assessed in wild-type eIF1A/*TIF11* or eIF1A/*tif11* R-to-A substitution mutant yeast strains expressing a plasmid with expression of eIF5^G31R^ (indicated by *sui5^-^*) or an empty vector (indicated by ev). Spot assays with ten-fold serial dilution (triangles) were performed using SD medium containing either 0.3 mM histidine (indicated as +HIS) or 0.003 mM histidine (indicated as -HIS). A) spot assay showing single substitution mutants of *tif11^R13A^, tif11^R14A^,* and *tif11^R62A^*; B) spot assay showing double substitution mutants of *tif11^R13,14A^, tif11^R13,62A^*, and *tif11^R14,62A^*; and C) spot assay showing triple substitution mutant *tif11^R13,14, 62A^*. Two biological replicates are shown from the same plate in each spot assay.

## Discussion

In our current work, we took advantage of a tailor-engineered protein arginine methyltransferase and a bioorthogonal profiling technique developed by the Luo group to comprehensively identify budding yeast *in vivo* methylated proteins by the major type I PRMT, Hmt1. In comparison with an antibody-based approach, our method identified 540 arginine methylated proteins – a significantly higher number of candidate substrates than any of the previously published work on Hmt1. A closer examination of our candidate methylated protein profile revealed an enrichment of regulators of translation initiation, among many gene-ontology categories. We characterized one such translation initiation factor, eIF1A, and demonstrated the importance of methylation in promoting translation initiation at near-cognate start codons.

Proteomic identification of arginine methylated proteins in cells poses a challenging task owing to the small size and lack of charge on methyl groups. Many existing approaches involve using antibodies specific to the methylarginines, which often recognizes flanking amino acids and thus possess limitations when used in unbiased screening experiments. Tracking arginine-methylated proteins to study their functional role in a cellular context requires high-quality reagents capable of differentiating their modification states, which are not always available. Approaches utilizing chemical biology to identify and characterize arginine-methylated proteins circumvent these limitations. One such approach relies on the use of a non-native cofactor recognized by a tailor-engineered PRMT of interest to modify substrates, resulting in substrates modified by a non-natural methylated arginine. This enables specific chemical functionality identification and tracking in a cellular context. However, many common trackable moieties, such as fluorescent molecules or short peptides for which antibodies exist commercially, are too large to fit in the active site of the PRMT of interest. Therefore, a strategy involving the installation of a small molecular handle capable of reacting selectively with an additionally introduced reactant attached to the reporter of interest allows for the formation of “click chemistry” reactions. This strategy pairs semi-synthetic PRMT substrate analogues containing clickable handles with PRMT mutants capable of accepting these substrates in an orthogonal manner to native substrates.

One candidate we identified from this profiling approach is the translation initiation factor eIF1A, which binds to the 40S subunit through interactions of its OB-fold with the body of the 40S ribosomal subunit, exerting control over start codon recognition through its C- and N-terminal tails that form alternating contacts with the PIC in either an open conformation, with tRNA in the P_out_ state, or a closed/P_in_ conformation, respectively (38, 44). The C-terminal tail (CTT) of eIF1A contains scanning enhancer (SE) elements that interact with the open state of the PIC with tRNA in the P_out_ state, while also hindering the conformational change to a closed state by physically occluding tRNA from occupying the P_in_, accommodated state (38). This open conformation of the 40S allows rapid and effective binding of tRNA and mRNA, and subsequent scanning of the complex. In contrast, the N-terminal tail (NTT) of eIF1A contains a scanning inhibitory element (SI_1_) and stabilizes the closed/P_in_ conformation of the PIC upon base-pairing, thereby arresting scanning and committing the ribosome to initiate translation at the codon present in the P site. Substitutions that compromise the C- or N-terminal tails promote or inhibit initiation at near-cognate start codons by stabilizing the closed or open conformation of the PIC, respectively (40, 41). Our translation fidelity results first established the involvement of Hmt1 in promoting initiation at near-cognate start codons (Fig. 5), and the use of eIF1A substitution mutants (Fig. 6) further supported a role of eIF1A methylation in decreasing initiation fidelity. It should be noted that the methyl-deficiency observed in the eIF1A substitution mutants may be attributed to either the absence of methylatable arginine in eIF1A or the potential disruption of the eIF1A interactions with Hmt1 and/or the ribosome caused by the substitution mutations. Utilizing the substitution mutants to demonstrate the functionality of R13 and R14 is not equivalent to having non-methylated arginine at these positions. Considering that Hmt1 methylates a wide range of substrates, isolating the specific contribution of methylation to the function of eIF1A in cells lacking Hmt1 presents a challenge. Future technical advancements that enable the specific removal of a modification without altering the amino acid context would provide a more optimal framework for defining the function of such modifications on a substrate. However, the similar effects of Hmt1 deletion and point mutations in eIF1A together support a role for Hmt1 methylation of eIF1A on start site selection.

While the *hmt1Δ* mutant did show increased fidelity, it did not display as strong of a ssu^-^ phenotype as the methylation-deficient point mutants of eIF1A (Figs. 5 and 6). It is possible that this difference in severity stems from disrupting the cumulative set of Hmt1 modifications on additional top candidate substrates identified, such as Ded1, Rps2, and Rps20, that influence initiation fidelity, which offsets the more severe effect observed by preventing individual eIF1A modifications. Ded1 (ortholog of human DDX3) is an essential DEAD-box RNA helicase that unwinds secondary structure to allow scanning of ribosomes, whereas Rps2 and Rps20 are two 40S proteins that are likely to affect movement of mRNA, PIC conformation, and start codon recognition (38, 45, 46). In addition to these high confidence substrates, components of the 43S complex known to affect start codon fidelity: eIF1/*SUI1*, eIF5/*TIF5*, two subunits of eIF2 (alpha/*SUI2,* gamma/*GCD11*) and four subunits of eIF3 (*PRT1, NIP1, TIF34, TIF35*); as well as components of the eIF4F helicase complex that affects ribosome loading and scanning (eIF4A/*TIF1*, and eIF4G/*TIF4631*) were also detected as putative Hmt1 substrates (Table S4), albeit at much lower modification levels than the aforementioned proteins. Modification of any of these initiation factors holds the potential to exert an opposing effect on start codon recognition that could dampen the effect of methylation of eIF1A and fine-tune start site recognition to an optimal level for balanced rates and fidelity of initiation. Alternatively, it is possible that methylation of additional substrates of Hmt1 outside of the translation initiation complex lead to an opposing effect on start codon recognition to dampen the ssu^-^ phenotype.

At the molecular level, methylation of these residues likely aids the eIF1A NTT in stabilizing the closed conformation of the PIC. Indeed, our substitution mutants at R13 and/or R14 in the critical NTT of eIF1A showed a strong ssu- phenotype, suggesting a decreased usage of the near-cognate start codon, consistent with reduced propensity to form the closed, P_in_ state. How would methylation at the R13 or R14 affect the function of eIF1A? Cryo-EM structures of yeast PICs in the closed/P_in_ state show that the NTT stabilizes the codon-anticodon helix of initiator tRNA and the start codon within the P site of the 40S. R13 and R14 residues are part of a disordered, likely flexible region that is not visible in the open state of the complex, but both residues are visible in the closed, P_in_ state structures. In structures without the N-terminus of eIF5 replacing eIF1, the R13 residue is in close proximity to threonine 3 and lysine 7 of eIF1A, and approaches an mRNA base, whereas the R14 residue faces rRNA and the lysine 11 sidechain of eIF1A (38). It is possible that methylation of R13 and R14 could promote the transition to the closed state by stabilizing interactions of the NTT with hydrophobic moieties of these potential contacts for a conformation that effectively interacts with the codon-anticodon helix. This would explain why non-methylatable alanine mutations of R13 and R14 affect start codon usage. In agreement with our results demonstrating that R13A and R14A substitutions can suppress the sui^-^ phenotype of the *sui5*^-^ mutation, it was previously reported that substitutions of these and three other basic residues of the eIF1A NTT found in uveal melanoma could suppress the sui- phenotype of the *SUI3-2* mutant of the eIF2 beta subunit (16). Combined with our findings, this further suggests that any effect of methylation on start codon recognition is exerted by stabilizing the closed/P_in_ conformation of the PIC, rather than anything more specific to *SUI3-2* or *sui5^-^*. In contrast, substitution mutation of R62, which resides in the OB fold of eIF1A, did not confer a phenotype in this assay, consistent with previous findings of eIF1A indicating that ssu^-^ mutations are primarily found in the N-terminal tail and helix alpha 2. Moreover, the methylated residues identified as being responsible for the bulk of the observed signal correlate with stronger phenotypes observed in the translational fidelity assay. Future experiments will help determine what role R62 methylation plays in eIF1A function that is not related to near-cognate start codon usage.

What is the biological significance behind the methylation at these residues on eIF1A? It is critical for cells to regulate their protein synthesis programs in response to environmental perturbations (47). Experimental evolution experiments have demonstrated that Hmt1-mediated methylation facilitates noise regulation in response to environmental stimuli, where adjustments in the heterogeneity of protein concentrations are made based on the cost and benefit of the proteins in question (47). In this study, it was determined that Hmt1 limits protein noise by modulating multiple pathways, but when under stress, Hmt1 levels are reduced and an increase in the heterogeneity of protein concentration is observed. It was revealed that this increase in the heterogeneity of protein concentration promotes cellular adaptation in response to such stress and hence, increases the evolutionary fitness of the organism (47). Though our results here demonstrate that a decrease in Hmt1-mediated methylation of eIF1A increases stringency in proper start codon recognition, and therefore less noise, it is possible that a need to have increased stringency for selective proteins during stress actually provides the efficiency and specificity required for such proteins. The transition from an open/P_out_ to closed/P_in_ state not only modulates the stringency of fidelity of AUG versus near-cognate start codon recognition but also mediates the ability of PICs to skip near-cognate codons and AUG codons in poor context. This phenomenon, termed leaky scanning, is probably best characterized as a mechanism of uORF skipping in the regulation of the general control pathway (48), whereby translation of several inhibitory uORFs is bypassed when eIF2 alpha is phosphorylated, allowing the production of the Gcn4 transcription factor. Recent work demonstrated that eIF1, by reducing the ability to recognize near-cognate start codons, limits translation of uORFs genome-wide in yeast. Our results suggest that methylation of eIF1A could normally serve the opposite purpose, to allow recognition of uORFs with near-cognate start codons or poor sequence context in response to stressors. It is possible that by allowing recognition of near-cognate uORFs, methylation of eIF1A prevents translation of the main ORFs of various proteins that stimulate alternative cellular programs under stress, such as Gcn4. By doing so, the decreased fidelity of start codon recognition may actually restrict heterogeneity in protein production. Under stress, reduced levels of Hmt1-mediated methylation could allow readthrough of particular uORFs instead, resulting in the production of alternative protein pools in the cell. For example, translation of housekeeping genes is often repressed under stress whereas translation of stress-induced transcripts can be maintained or even enhanced (49). Our data points to a possible role for eIF1A methylation in such a process. Further work is needed to determine the extent of Hmt1-mediated methylation on translation fidelity during stress.

## Acknowledgement

The authors thank Alan Hinnebusch and Jon Lorsch for providing plasmids and strains used in this study, Tanweer Hussain for commenting on structural effects of eIF1A methylation on 43S interactions, and members of the Yu, Walker, and Luo laboratories for helpful discussions. This work was supported by an NSF award (MCB-2100563) to M.C.Y., NIH grants R00GM119173 and R01GM139977 and UB CAS start-up funds to S.W., and NIH grants R35GM131858 and P30CA008748 to M.L.

## Supplemental data

**Table 1. Table of Yeast Strains Used in this Study.**

**Table 2. Table of Primers Used in this Study.**

**Table 3. Table of Plasmids Used in this Study.**

**Table 4. List of Candidate Proteins Identified from Bioorthogonal Profiling**

## References

1. Bedford, M. T., and Clarke, S. G. (2009) Protein arginine methylation in mammals: who, what, and why. Molecular cell 33, 1–13

2. Xu, J., and Richard, S. (2021) Cellular pathways influenced by protein arginine methylation: Implications for cancer. Mol Cell 81, 4357–4368

3. Lorton, B. M., and Shechter, D. (2019) Cellular consequences of arginine methylation. Cell Mol Life Sci 76, 2933–2956

4. Gary, J. D., Lin, W.-J., Yang, M. C., Herschman, H. R., and Clarke, S. (1996) The predominant protein-arginine methyltransferase from Saccharomyces cerevisiae. Journal of Biological Chemistry 271, 12585–12594

5. Tang, J., Frankel, A., Cook, R. J., Kim, S., Paik, W. K., Williams, K. R., Clarke, S., and Herschman, H. R. (2000) PRMT1 is the predominant type I protein arginine methyltransferase in mammalian cells. Journal of Biological Chemistry 275, 7723–7730

6. Hamey, J. J., Separovich, R. J., and Wilkins, M. R. (2018) MT-MAMS: Protein Methyltransferase Motif Analysis by Mass Spectrometry. J Proteome Res 17, 3485–3491

7. Jackson, C. A., Yadav, N., Min, S., Li, J., Milliman, E. J., Qu, J., Chen, Y.-C., and Yu, M. C. (2012) Proteomic analysis of interactors for yeast protein arginine methyltransferase hmt1 reveals novel substrate and insights into additional biological roles. Proteomics 12, 3304–3314

8. Low, J. K., Im, H., Erce, M. A., Hart-Smith, G., Snyder, M. P., and Wilkins, M. R. (2016) Protein substrates of the arginine methyltransferase Hmt1 identified by proteome arrays. Proteomics 16, 465–476

9. Yagoub, D., Hart-Smith, G., Moecking, J., Erce, M. A., and Wilkins, M. R. (2015) Yeast proteins Gar1p, Nop1p, Npl3p, Nsr1p, and Rps2p are natively methylated and are substrates of the arginine methyltransferase Hmt1p. Proteomics 15, 3209–3218

10. Hartel, N. G., Chew, B., Qin, J., Xu, J., and Graham, N. A. (2019) Deep Protein Methylation Profiling by Combined Chemical and Immunoaffinity Approaches Reveals Novel PRMT1 Targets. Mol Cell Proteomics 18, 2149–2164

11. Luo, M. (2012) Current chemical biology approaches to interrogate protein methyltransferases. ACS chemical biology 7, 443–463

12. Blum, G., Islam, K., and Luo, M. (2013) Bioorthogonal Profiling of Protein Methylation (BPPM) Using an Azido Analog of S-Adenosyl-L-Methionine. Current protocols in chemical biology 5, 45–66

13. Islam, K., Chen, Y., Wu, H., Bothwell, I. R., Blum, G. J., Zeng, H., Dong, A., Zheng, W., Min, J., Deng, H., and Luo, M. (2013) Defining efficient enzyme-cofactor pairs for bioorthogonal profiling of protein methylation. Proceedings of the National Academy of Sciences of the United States of America 110, 16778–16783

14. Polevoda, B., and Sherman, F. (2007) Methylation of proteins involved in translation. Molecular microbiology 65, 590–606

15. Haghandish, N., Baldwin, R. M., Morettin, A., Dawit, H. T., Adhikary, H., Masson, J.-Y., Mazroui, R., Trinkle-Mulcahy, L., and Côté, J. (2019) PRMT7 methylates eukaryotic translation initiation factor 2α and regulates its role in stress granule formation. Molecular Biology of the Cell 30, 778–793

16. Martin-Marcos, P., Zhou, F., Karunasiri, C., Zhang, F., Dong, J., Nanda, J., Kulkarni, S. D., Sen, N. D., Tamame, M., Zeschnigk, M., Lorsch, J. R., and Hinnebusch, A. G. (2017) eIF1A residues implicated in cancer stabilize translation preinitiation complexes and favor suboptimal initiation sites in yeast. Elife 6

17. Kozak, M. (1978) How do eucaryotic ribosomes select initiation regions in messenger RNA? Cell 15, 1109–1123

18. Furic, L., Rong, L., Larsson, O., Koumakpayi, I. H., Yoshida, K., Brueschke, A., Petroulakis, E., Robichaud, N., Pollak, M., and Gaboury, L. A. (2010) eIF4E phosphorylation promotes tumorigenesis and is associated with prostate cancer progression. Proceedings of the National Academy of Sciences 107, 14134–14139

19. Clemens, M. J. (2001) Initiation factor eIF2α phosphorylation in stress responses and apoptosis. Signaling Pathways for Translation, pp. 57–89, Springer

20. Knop, M., Siegers, K., Pereira, G., Zachariae, W., Winsor, B., Nasmyth, K., and Schiebel, E. (1999) Epitope tagging of yeast genes using a PCR-based strategy: more tags and improved practical routines. Yeast (Chichester, England) 15, 963–972

21. Longtine, M. S., Mckenzie III, A., Demarini, D. J., Shah, N. G., Wach, A., Brachat, A., Philippsen, P., and Pringle, J. R. (1998) Additional modules for versatile and economical PCR-based gene deletion and modification in Saccharomyces cerevisiae. Yeast 14, 953–961

22. Lee, B., Udagawa, T., Singh, C. R., and Asano, K. (2007) Yeast phenotypic assays on translational control. Methods in enzymology, pp. 105–137, Elsevier

23. Jackson, C. A., and Michael, C. Y. (2014) Detection of protein arginine methylation in Saccharomyces cerevisiae. Yeast Protocols, pp. 229–247, Springer

24. Acker, M. G., Kolitz, S. E., Mitchell, S. F., Nanda, J. S., and Lorsch, J. R. (2007) Reconstitution of yeast translation initiation. Methods in enzymology, pp. 111–145, Elsevier

25. Gietz, R. D., and Sugino, A. (1988) New yeast-Escherichia coli shuttle vectors constructed with in vitro mutagenized yeast genes lacking six-base pair restriction sites. Gene 74, 527–534

26. Wang, R., Zheng, W., Yu, H., Deng, H., and Luo, M. (2011) Labeling substrates of protein arginine methyltransferase with engineered enzymes and matched S-adenosyl-L-methionine analogues. Journal of the American Chemical Society 133, 7648–7651

27. Henry, M. F., and Silver, P. A. (1996) A novel methyltransferase (Hmt1p) modifies poly(A)+-RNA-binding proteins. Molecular and cellular biology 16, 3668–3678

28. Siebel, C. W., and Guthrie, C. (1996) The essential yeast RNA binding protein Np13p is methylated. Proceedings of the National Academy of Sciences of the United States of America 93, 13641–13646

29. McBride, A. E., Weiss, V. H., Kim, H. K., Hogle, J. M., and Silver, P. A. (2000) Analysis of the yeast arginine methyltransferase Hmt1p/Rmt1p and its in vivo function. Cofactor binding and substrate interactions. The Journal of biological chemistry 275, 3128–3136

30. Pestova, T. V., Borukhov, S. I., and Hellen, C. U. (1998) Eukaryotic ribosomes require initiation factors 1 and 1A to locate initiation codons. Nature 394, 854–859

31. Acker, M. G., Kolitz, S. E., Mitchell, S. F., Nanda, J. S., and Lorsch, J. R. (2007) Reconstitution of yeast translation initiation. Methods in enzymology 430, 111–145

32. Wooderchak, W. L., Zang, T., Zhou, Z. S., Acuna, M., Tahara, S. M., and Hevel, J. M. (2008) Substrate profiling of PRMT1 reveals amino acid sequences that extend beyond the “RGG” paradigm. Biochemistry 47, 9456–9466

33. Gary, J. D., and Clarke, S. (1998) RNA and protein interactions modulated by protein arginine methylation. Progress in nucleic acid research and molecular biology, pp. 65–131, Elsevier

34. Maag, D., Fekete, C. A., Gryczynski, Z., and Lorsch, J. R. (2005) A conformational change in the eukaryotic translation preinitiation complex and release of eIF1 signal recognition of the start codon. Molecular cell 17, 265–275

35. Passmore, L. A., Schmeing, T. M., Maag, D., Applefield, D. J., Acker, M. G., Algire, M. A., Lorsch, J. R., and Ramakrishnan, V. (2007) The eukaryotic translation initiation factors eIF1 and eIF1A induce an open conformation of the 40S ribosome. Molecular cell 26, 41–50

36. Yu, Y., Marintchev, A., Kolupaeva, V. G., Unbehaun, A., Veryasova, T., Lai, S. C., Hong, P., Wagner, G., Hellen, C. U., and Pestova, T. V. (2009) Position of eukaryotic translation initiation factor eIF1A on the 40S ribosomal subunit mapped by directed hydroxyl radical probing. Nucleic acids research 37, 5167–5182

37. Weisser, M., Voigts-Hoffmann, F., Rabl, J., Leibundgut, M., and Ban, N. (2013) The crystal structure of the eukaryotic 40S ribosomal subunit in complex with eIF1 and eIF1A. Nature structural & molecular biology 20, 1015–1017

38. Hussain, T., Llácer, J. L., Fernández, I. S., Munoz, A., Martin-Marcos, P., Savva, C. G., Lorsch, J. R., Hinnebusch, A. G., and Ramakrishnan, V. (2014) Structural changes enable start codon recognition by the eukaryotic translation initiation complex. Cell 159, 597–607

39. Fekete, C. A., Applefield, D. J., Blakely, S. A., Shirokikh, N., Pestova, T., Lorsch, J. R., and Hinnebusch, A. G. (2005) The eIF1A C-terminal domain promotes initiation complex assembly, scanning and AUG selection in vivo. The EMBO journal 24, 3588–3601

40. Fekete, C. A., Mitchell, S. F., Cherkasova, V. A., Applefield, D., Algire, M. A., Maag, D., Saini, A. K., Lorsch, J. R., and Hinnebusch, A. G. (2007) N-and C-terminal residues of eIF1A have opposing effects on the fidelity of start codon selection. The EMBO journal 26, 1602–1614

41. Saini, A. K., Nanda, J. S., Lorsch, J. R., and Hinnebusch, A. G. (2010) Regulatory elements in eIF1A control the fidelity of start codon selection by modulating tRNAiMet binding to the ribosome. Genes & development 24, 97–110

42. Nanda, J. S., Saini, A. K., Munoz, A. M., Hinnebusch, A. G., and Lorsch, J. R. (2013) Coordinated movements of eukaryotic translation initiation factors eIF1, eIF1A, and eIF5 trigger phosphate release from eIF2 in response to start codon recognition by the ribosomal preinitiation complex. The Journal of biological chemistry 288, 5316–5329

43. Saini, A. K., Nanda, J. S., Martin-Marcos, P., Dong, J., Zhang, F., Bhardwaj, M., Lorsch, J. R., and Hinnebusch, A. G. (2014) Eukaryotic translation initiation factor eIF5 promotes the accuracy of start codon recognition by regulating Pi release and conformational transitions of the preinitiation complex. Nucleic acids research 42, 9623–9640

44. de Breyne, S., Yu, Y., Unbehaun, A., Pestova, T. V., and Hellen, C. U. (2009) Direct functional interaction of initiation factor eIF4G with type 1 internal ribosomal entry sites. Proceedings of the National Academy of Sciences of the United States of America 106, 9197–9202

45. Sen, N. D., Zhou, F., Harris, M. S., Ingolia, N. T., and Hinnebusch, A. G. (2016) eIF4B stimulates translation of long mRNAs with structured 5’ UTRs and low closed-loop potential but weak dependence on eIF4G. Proceedings of the National Academy of Sciences of the United States of America 113, 10464–10472

46. Walker, S. E., Zhou, F., Mitchell, S. F., Larson, V. S., Valasek, L., Hinnebusch, A. G., and Lorsch, J. R. (2013) Yeast eIF4B binds to the head of the 40S ribosomal subunit and promotes mRNA recruitment through its N-terminal and internal repeat domains. Rna 19, 191–207

47. You, S.-T., Jhou, Y.-T., Kao, C.-F., and Leu, J.-Y. (2019) Experimental evolution reveals a general role for the methyltransferase Hmt1 in noise buffering. PLoS biology 17

48. Hinnebusch, A. G. (2005) Translational regulation of GCN4 and the general amino acid control of yeast. Annu. Rev. Microbiol. 59, 407–450

49. Anderson, P., and Kedersha, N. (2002) Stressful initiations. Journal of cell science 115, 3227–3234

